# Gene signatures and host-parasite interactions revealed by dual single-cell profiling of *Plasmodium vivax* liver infection

**DOI:** 10.1101/2021.09.01.454555

**Authors:** Liliana Mancio-Silva, Nil Gural, Marc H. Wadsworth, Vincent L. Butty, Travis K. Hughes, Sandra March, Niketa Nerurkar, Wanlapa Roobsoong, Heather E. Fleming, Charlie Whittaker, Stuart S. Levine, Jetsumon Sattabongkot, Alex K. Shalek, Sangeeta N. Bhatia

**Affiliations:** Institute for Medical Engineering and Science, MIT, Cambridge, MA, 02142 USA; David H. Koch Institute for Integrative Cancer Research, MIT, Cambridge, MA, 02142 USA; Institut National de la Santé et de la Recherche Médicale, Unité 1201, Paris, France; Department of Chemistry, MIT, Cambridge, MA 02142, USA; BioMicro Center, MIT, Cambridge, MA 02142, USA; Mahidol Vivax Research Unit, Faculty of Tropical Medicine Mahidol University, Bangkok 10400, Thailand; Ragon Institute of MGH, MIT, and Harvard, Cambridge, MA 02142, USA; Broad Institute of MIT and Harvard, Cambridge, MA 02142, USA; Howard Hughes Medical Institute, Chevy Chase, MD, 20815 USA

**Keywords:** *Plasmodium vivax*, hypnozoites, dormancy, hepatocytes, single-cell, transcriptomics, host-parasite interactions

## Abstract

Malaria-causing *P. vivax* parasites can linger in the human liver for weeks to years, and then reactivate to cause recurrent blood-stage infection. While an important target for malaria eradication, little is known about the molecular features of the replicative and non-replicative states of intracellular *P. vivax* parasites, or the human host-cell responses to them. Here, we leverage a bioengineered human microliver platform to culture Thai clinical isolates of *P. vivax* in primary human hepatocytes and conduct transcriptional profiling of infected cultures. By coupling enrichment strategies with bulk and single-cell analyses, we captured both parasite and host transcripts in individual hepatocytes throughout the infection course. We defined host- and state-dependent transcriptional signatures and identified previously unappreciated populations of replicative and non-replicative parasites, sharing features with sexual transmissive forms. We found that infection suppresses transcription of key hepatocyte function genes, and that *P. vivax* elicits an innate immune response that can be manipulated to control infection. Our work provides an extendible framework and resource for understanding host-parasite interactions and reveals new insights into the biology of malaria dormancy and transmission.

## INTRODUCTION

*Plasmodium* parasite, the causative pathogen of malaria, has a complex life cycle that spans multiple hosts. Disease transmission is initiated upon bite of an infected *Anopheles* mosquito, which deposits infective parasites, called sporozoites, into the human host. Sporozoites travel to the liver, invade hepatocytes and replicate, forming thousands of new parasites called merozoites, which eventually break out into the blood stream, cyclically invading erythrocytes and initiating clinical symptoms. Parasite transmission back to the mosquito is ensured by the development of sexual gametocyte forms during the asexual erythrocytic cycle that are taken up upon bite to restart the life cycle in the insect host. Uniquely, in the case of *P. vivax*, a subset of liver-stage parasites develops into dormant forms called hypnozoites, which can re-activate weeks to years after initial infection to cause relapsing disease. Thus, the liver-stage, which is obligate yet clinically silent and includes relapse-causing hypnozoites, presents a unique opportunity for malaria intervention before onset of symptoms. However, our knowledge of liver-stage malaria, and the response of its hepatocyte host is sparse due to difficult access to the parasite and lack of suitable human liver models. To date, much of our historical knowledge has been based on liver biopsies of infected patients, making it challenging to perform mechanistic studies on liver-stage forms, especially the quiescent hypnozoites. Transcriptomic studies hold promise for unveiling mechanistic insight into liver-stage *P. vivax* relapsing biology, but the low infection rate and the reduced quantity of parasite transcripts in a transcriptionally active host cell environment has made it difficult to perform these studies.

We recently provided some of the first insights into the transcriptional features of *P. vivax* liver-stages by leveraging an *in vitro* primary human liver platform (MPCC, micropatterned co-cultures) that recapitulates key aspects of *P. vivax* liver-stage biology, including establishment of persistent dormant forms, growing schizonts, merozoite release, and subsequent infection of overlaid erythrocytes. Our work revealed reduced transcriptional activity in hypnozoite-enriched samples, specifically, suppressed transcripts for functions related to cell division and invasion machinery, consistent with a quiescent state (Gural et al., 2018). However, the single time-point bulk sequencing used in this study prevented us to capture the inherent heterogeneity of the distinct liver-stage parasite forms. Achieving a deeper understanding of pathogen-host interactions has the potential to provide insight into mechanisms that could be leveraged to treat or prevent infection, as suggested by innate interferon responses to infection by rodent malaria parasites (Liehl et al., 2013) and a number of viruses (Schneider et al., 2014). However, a closer look into infection-specific host responses and potential protective responses in uninfected neighbors requires single-cell resolution. Recently, diverse single-cell technologies have revealed stage-specific transcriptional signatures in mosquito (Real et al., 2021), blood and gametocyte (Poran et al., 2017; Walzer et al., 2018) stages from non-relapsing human parasites, and the entire life cycle of rodent parasites (Howick et al., 2019), all of which can be easily cultured and propagated in laboratories. For *P. vivax*, single-cell transcriptomic studies have been conducted in blood-stages collected from infected monkeys (Sà et al., 2020), but analysis of liver infection has not been performed to date.

Here, we present the first comprehensive view of the liver-stage transcriptomes of a human-infecting malaria parasite and its host cells at single-cell resolution. To achieve this goal, we coupled MPCC with Seq-Well, a recently developed low-cost and portable single-cell platform that does not require fluorescent labeling and is compatible with use of samples collected in endemic settings (Gierahn et al., 2017; Hughes et al., 2020). With this combined platform, we define distinct signatures between early, dormant, mid, and late-stage parasites, and identify a sub-population of sexually committed forms in the liver, previously thought to appear only during erythrocytic infection. We interrogate pathogen-host interactions and describe innate immune responses by uninfected bystander cells, representing a likely mechanism for endogenous protection from secondary infections. We validate expression of host candidates and report cytokine- and stage-dependent anti-parasite activity by induction of interferon signaling. Together, the data presented here provide a closer look at transcriptional signatures in *P. vivax* liver-stages, including host responses, and offer novel insights into their unique biology.

## RESULTS

### Profiling liver-stage *P. vivax* infection by pairing targeted sequencing with single-cell analysis

To comprehensively profile *P. vivax* liver-stage infection, we collected cells from MPCC cultures of primary human hepatocytes at multiple time points following infection (**Figures 1A-B, S1A)**. Sampling spanned the full liver-stage developmental period (days 1 to 11), comprising a mix of both replicating schizonts and non-replicative hypnozoites. To obtain hypnozoite-enriched samples, MPCCs were treated from day 5 to 8 with a phosphatidylinositol 4-kinase (Pi4K) inhibitor compound, a dosing regimen that eliminates the replicative parasites while preserving the dormant parasites (Gural et al., 2018). For hypnozoite-enrichment without drug treatment, we collected cultures at day 14. To obtain a baseline reading of the host response, naïve and mock-exposed MPCCs were prepared and collected in parallel. A total of 56 samples from two independent infections performed with clinical Thai *P. vivax* isolates were processed for high throughput single-cell RNA sequencing (scRNA-seq) using Seq-Well S^3^ (Hughes et al., 2020). Samples were also collected in bulk for RNA sequencing (RNA-seq) analysis (**Figure 1A**).

**Figure 1.**
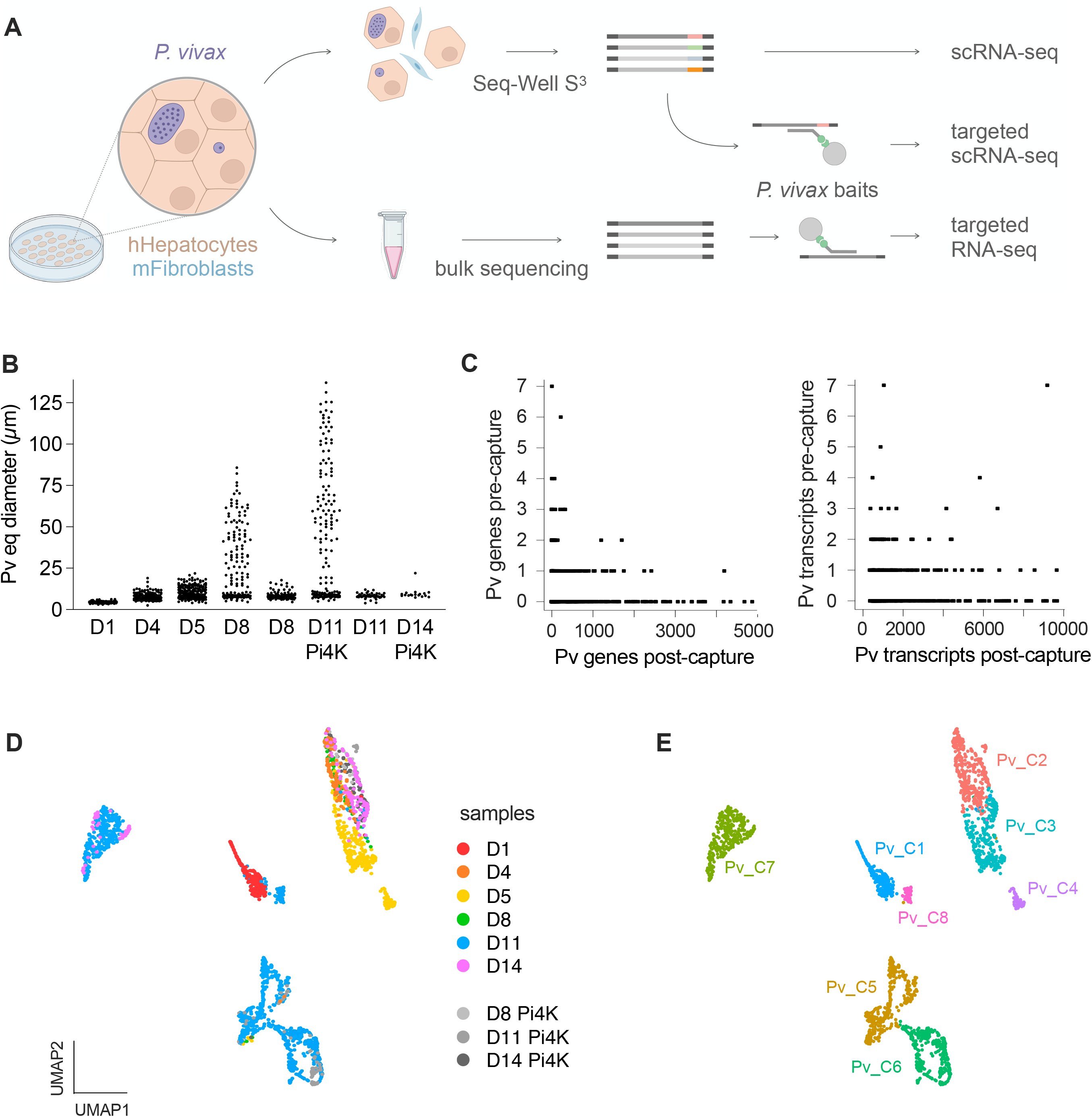
Single-cell profiling of *P. vivax* liver-stage infection. **A.** Illustration of *P. vivax* liver-stages (purple) maintained in micropatterned co-cultures (MPCC) of human hepatocytes (brown) and supportive mouse fibroblasts (blue). Cultures were dissociated with trypsin (top) or directly collected in bulk (bottom). Sample processing pipeline is shown for whole transcriptome (top) and targeted (middle) single-cell and targeted bulk RNA sequencing (bottom). **B.** Parasite size distribution and collection days. Day 14 and PiK4-treated samples are enriched in nonreplicative hypnozoites (parasites < 10 μm diameter). Each dot represents an individual parasite (*n* = 3-8 wells pooled from 2 independent infections). Quantification of parasite numbers given in **Figure S1A**. **C.** Scatterplots comparing the number of *P. vivax* genes (left) and transcripts (right) prior and post-capture with parasite-specific baits. Efficient depletion of host genes shown in **Figure S1B**. **D-E.** UMAP of 1,991 individual *P. vivax* parasites colored by sample identity (**D**) and parasite cluster type (**E**) using Scanorama. Heatmap showing cluster-specific marker genes provided in **Figure S2A**.

Because whole transcriptome scRNA-seq resulted in poor representation of *P. vivax* genes and transcripts, we incorporated an additional step whereby barcoded parasite transcripts were enriched using previously validated nucleic acid baits targeting the entire *P. vivax* genome (Gural et al., 2018). Capture and re-sequencing of samples increased the efficiency of *P. vivax* gene and transcript detection by 5.6-fold per single parasite (**Figures 1C, S1B**). Combining single-cell transcriptomes from pre- and post-capture, we recovered 1,991 individual parasites with greater than 10 genes or 100 *P. vivax*-mapped transcripts. This tally fairly reflects the number of hepatic infections at the different timepoints, except for day 11 samples from which a higher number of parasites was recovered (likely representing free merozoites released from mature schizonts during sample processing; **Figure S1A; Table S1**).

### Single-cell profiling *P. vivax* liver infection defines stage-specific gene signatures

Integration of parasite transcriptomes with Scanorama yielded 8 *P. vivax* clusters, corresponding to distinct developmental liver-stages that were visualized by uniform manifold approximation and projection (UMAP; **Figures 1D-E**). Cluster Pv_C1 contains early-stage individual parasites (day 1), while clusters Pv_C2-C4 comprise *P. vivax* parasites present in the mixed samples collected on days 4 to 8, as well the hypnozoites from Pi4K-enriched cultures and the day 14 samples. Late-stage schizont parasites (day 11) are scattered across clusters Pv_C5-C8.

Based on the gene patterns that define each cluster (**Figure S2A; Table S1**), early liver-stage parasites (Pv_C1) are characterized by residual expression of sporozoite-specific genes and expression of genes involved in cytoskeleton organization process, likely necessary to establish intracellular infection. Pv_C2-C4 clusters appear to represent a core mid-stage liver program, comprising genes implicated in housekeeping functions important for parasite growth such as translation (*EIF1D,G; EIF2A,B; EIF3A-E*) and metabolism (*ENO*, *ACC, HCS1*). Well-characterized genes such as those encoding liver-specific protein 1 (*LISP1*) and 2 (*LISP2*) appear as top marker genes for these clusters. Progressing along the liver-stage development, GO terms for cell cycle, mitotic division, and adhesion of symbiont to host become significantly represented in Pv_C5-C6. Gene cluster identifiers include several members of merozoite surface proteins (*MSP*), serine repeat antigen (*SERA*), rhoptry associated (*RAP, RAMA*) and rhoptry neck (*RON*) multigene families that are necessary for red blood cell invasion, and also the merozoite egress subtilisin-like protease 1 (*SUB1*). Finally, among the late-stage parasites Pv_C7-C8 represent a previously unidentified population of hepatic parasites that co-express merozoite- and gametocyte-specific genes. Marker genes for these clusters include multiple copies of the tryptophan-rich protein family, which are expressed by merozoites of multiple *Plasmodium* species and have been implicated in erythrocyte invasion, the gametocyte antigen *G27/25,* the gamete release protein (*GAMER*) and the homolog of gametocyte exported protein 5 (*GEXP5*, aka *PHISTc*).

### Transcriptional profiling of *P. vivax* liver-stages reveals early sexual commitment

To gain further insight into the subpopulation of parasites defined by clusters Pv_C7 and Pv_C8, we searched the dataset for additional gametocyte-specific genes. Expression of canonical sexual markers (*PVS16, P28, P230),* including male (*MGET* and *MDG1*) and female (*RUBV1* and *G377*) specific genes recently assigned to *P. vivax* (Sà et al., 2020), was detected throughout the developmental liver-stages in multiple clusters (**Figure 2A**). Their expression was also confirmed via targeted bulk RNA-seq (**Figure 2B; Table S2**) and by RT-qPCR analysis of a small subset of genes (**Figure 2C**). For further validation, we performed *in situ* hybridization for *GEXP5*, which appears highly expressed in day 11 samples (**Figures 2B, 2C**) and is known as an early sexual stage marker in blood-stages of the human malaria parasite *P. falciparum* (Tibúrcio et al., 2015). We found *GEXP5* transcripts in 5-15% of schizonts, with positive signal in all merozoites, while the remaining schizonts did not contain this transcript (**Figure 2D**). Interestingly, we found that 16-25% of parasites in clusters Pv_C1 and Pv_C6 (**Figure 2A; Table S1**) express the gene encoding AP2-G, the transcriptional master regulator of sexual development in blood stages (Kafsack et al., 2014; Sinha et al., 2014). Exploring the dynamics of this transcription factor, we found induced expression of *AP2-G* and its upstream activator, gametocyte development 1 (*GDV1*) (Filarsky et al., 2018), as early as day 1 after hepatocyte invasion. Expression of *PVS16*, a downstream target of AP2-G increased from day 2, and onwards (**Figure 2E**). PVS16 protein has been detected in a fraction of *P. vivax* hepatic schizonts at day 8 (Roth et al., 2018; Schafer et al., 2020). Altogether, the data indicates that commitment to gametocytogenesis might occur early during liver-stage development in a subset of parasites, likely leading to formation of sexually committed merozoites.

**Figure 2.**
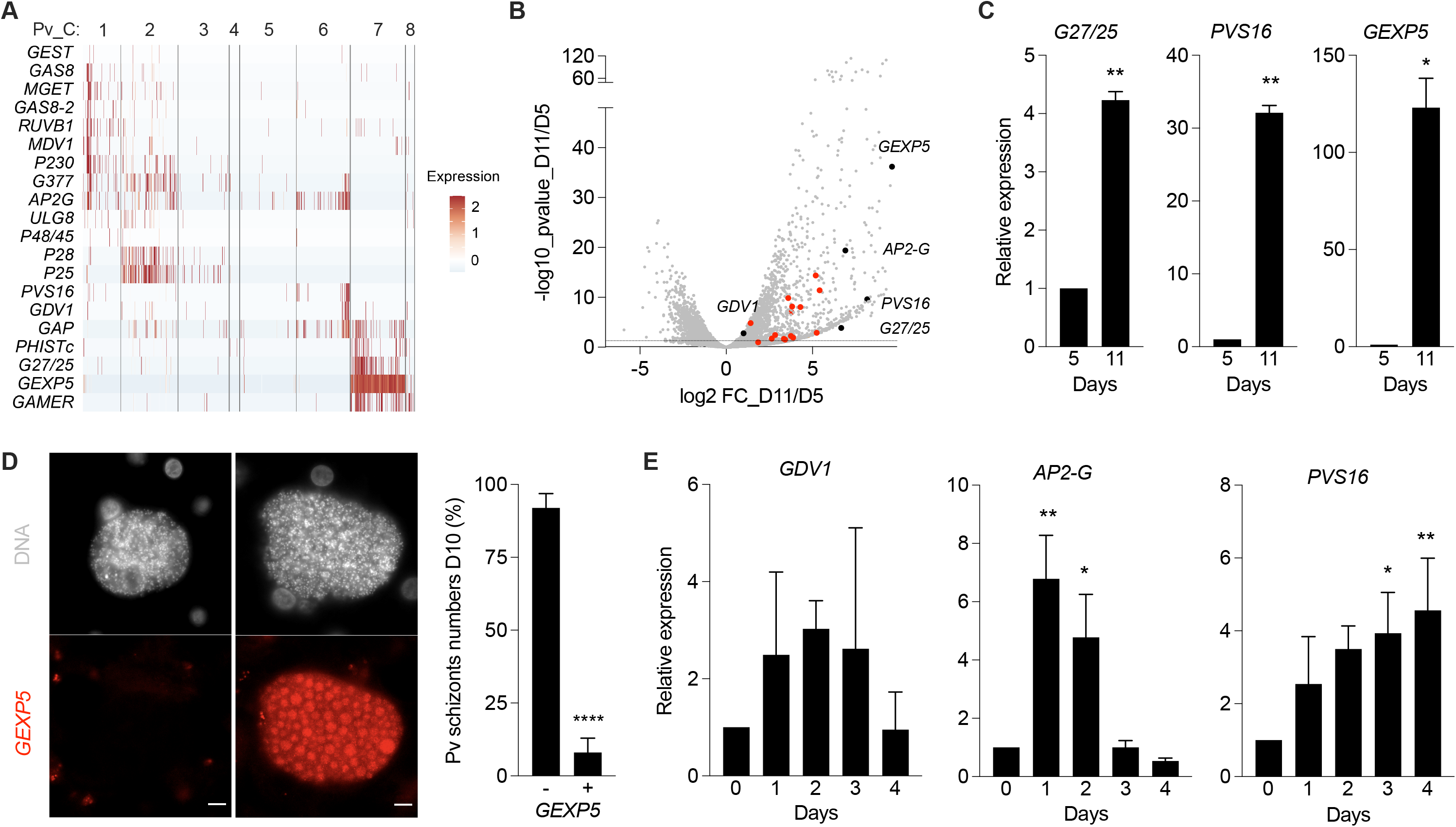
Characterization of *P. vivax* individual transcriptomes. **A.** Heatmap showing the expression of gametocyte-specific genes for each parasite cluster. 100 parasites are shown, except clusters Pv_C4 and Pv_C8 that contain fewer cells. Each bar represents a single parasite. **B.** Volcano plot comparing differential expression of day 5 and day 11 samples on targeted bulk RNA-seq. Parasite gametocyte gene markers highlighted. Red for gametocyte-genes described in (Sà et al., 2020). Black for genes validated in **C-E**. **C.** Relative expression of gametocyte markers by quantitative RT-PCR (mean ± SEM; *n* = 2 independent infections; *t*-test: *, *p* < 0.05; **, *p* < 0.01). **D.** RNA in situ hybridization of *P. vivax* parasites at day 10. *GEXP5* transcripts shown in red. Scale bars, 10 μm. Quantification of *GEXP5*-positive and -negative schizonts (mean ± SEM; *n* = 4 wells; *t*-test: ****, *p* < 0.0001). **E.** Relative expression of *GDV1, AP2-G* and *PVS16* by quantitative RT-PCR at early timepoints (mean ± SEM; *n* = 2 independent infections; 1-way ANOVA test: *, *p* < 0.05; **, *p* < 0.01). FC, fold chnage; *G27/25,* gametocyte antigen; *PVS16,* parasitophorous vacuole membrane protein S16; *GAMER*, gamete release protein; *GEXP5*, gametocyte exported protein 5; *GDV1*, gametocyte development 1; *AP2-G*, AP2 domain transcription factor regulator of gametocytogenesis.

### *P. vivax* non-replicative liver-stages depend on proteolytic activity and are sexually committed

To inform the identification of hypnozoite-specific gene signature(s), we re-clustered the mid-stage parasite clusters Pv_C2-C4 (**Figures 3A-B; Table S1**). This enabled us to explore transcriptional differences among the mixed and hypnozoite-containing samples and revealed a sub-cluster (Pv_SC3) significantly enriched in genes encoding proteins with peptidase activity (10 genes) and nucleic acid binding proprieties (18 genes) in the non-replicative parasite population. These include several members of protease families (vivapains and plasmepsins), members of ApiAP2 family of transcription factors (*AP2-Tel* and *AP2-FG*) and *PUF1,* a Pumilio RNA binding protein known to be involved in translational repression (Bennink and Pradel, 2019). Consistent with a quiescent state, parasites in Pv_SC3 show low levels of the replicative marker *LISP2* (**Figure 3C**). The expression of transcriptional regulators of gametocytogenesis *AP2-G* and *AP2-FG* prompted us to look for additional gametocyte-related genes (as in **Figure 2A**). Remarkably, we found expression of canonical sexual markers (*P28, P25, P230*) in Pv_SC3 subcluster, suggesting that a subpopulation of *P. vivax* quiescent hypnozoites could be pre-committed to become sexually transmissive forms (**Figures 3C-D**). A larger number of gametocyte-specific genes however was found in the Pv_SC5 subcluster, which is largely *LISP2* positive. Thus, Pi4K-treated and day 14 parasites in Pv_SC5 subcluster likely represent a subpopulation of sexually committed reactivating hypnozoites. Presence of gametocyte-specific transcripts in Pi4K-treated samples was confirmed by RT-qPCR analysis, further supporting the concept that non-replicative parasites could be sexually committed (**Figure S3A**).

**Figure 3.**
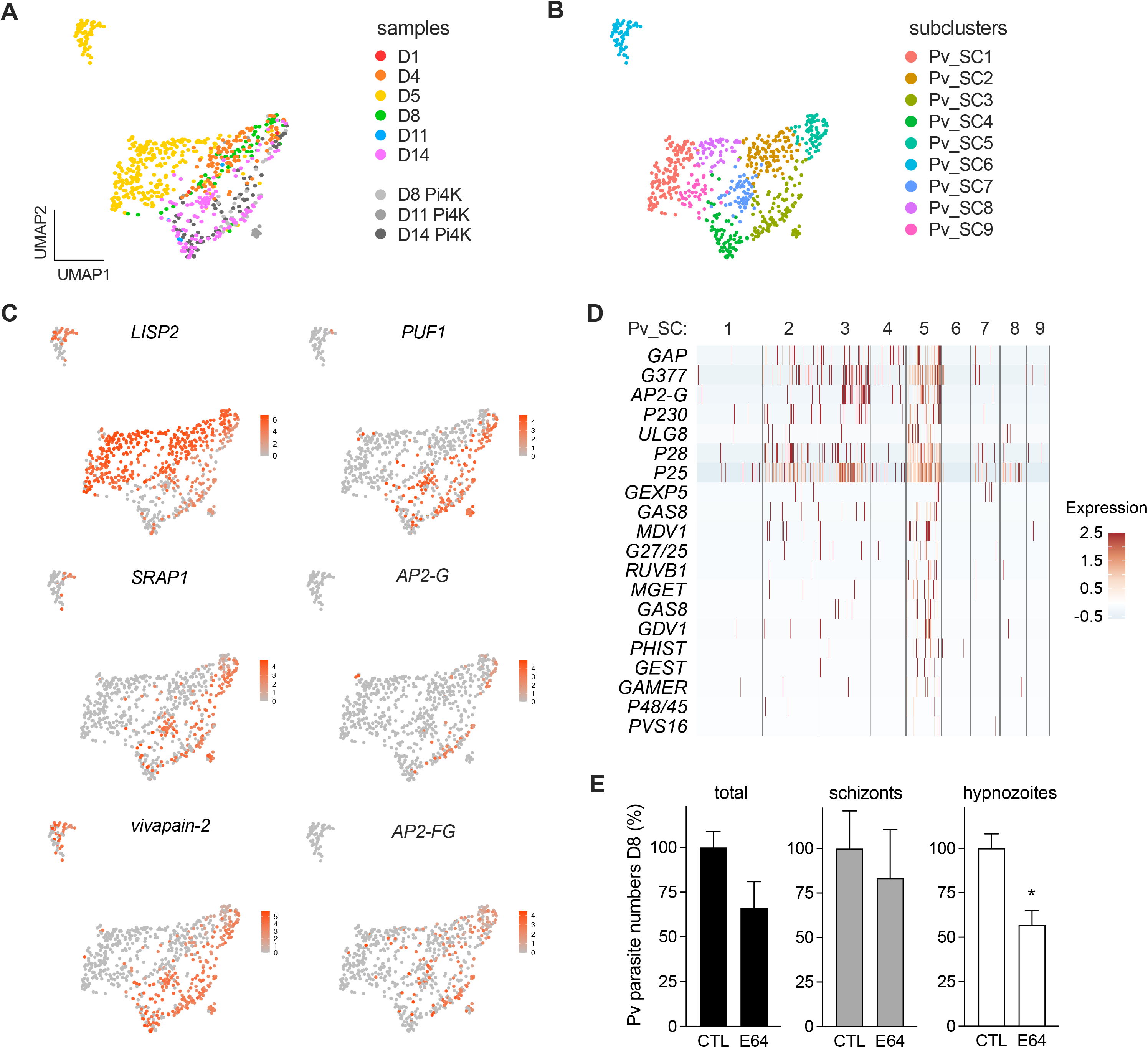
Characterization of *P. vivax* mid-stage transcriptomes. **A-B.** UMAP of 713 *P. vivax* parasites (Pv_C2-C4 re-clustered) colored by sample identity (**A**) and parasite subcluster types (**B**). **C.** UMAP as in **A** highlighting expression levels of *LISP2*, *PUF1*, genes encoding peptidases (*SRAP1* and *PVP01_0916200 vivapain-2*) and ApiAP2s (*AP2-G* and *AP2-FG*). **D.** Heatmap showing expression of gametocyte-specific genes for each parasite subcluster. Gene list as in **Figure 2A**. Each bar represents a single parasite. Expression of gametocyte markers in Pi4K-treated samples is shown in **Figure S3A**. **E.** Treatment (days 0-8) of *P. vivax* infected cultures with protease inhibitor E64 (1 μM; mean ± SEM; *n* = 3-5 wells; *t* -test: *, *p* < 0.05). Parasite size distribution is given in **Figure S3B**. *LISP2*, liver-specific protein 2; *PUF1*, Pumilio RNA binding p rotein; *SRAP1*, sea star regeneration associated protease; *AP2-G*, transcription factor regulator of gametocytogenesis; A *P2-FG*, female gametocyte-specific t ranscriptional regulator.

Protein degradation is important for maintenance of a hibernatory state induced by nutrient limitation during the *P. falciparum* erythrocytic cycle (Babbitt et al., 2012). To determine whether proteolytic activity could be required to maintain viability of quiescent hypnozoites, we exposed *P. vivax*-infected MPCCs to E64, a membrane-permeable protease inhibitor. No effect was found on schizont forms, however, the number of *P. vivax* hypnozoites was significantly reduced in the E64-treated cultures (**Figures 3E, S3B**). Altogether the results suggest that *P. vivax* hypnozoites in the liver could represent a reservoir of sexually committed parasites and dormancy might depend on transcription and translational repression mechanisms and proteolysis for long-lasting viability.

### Pre-erythrocytic sexual differentiation is associated to distinct host metabolic states

To characterize host cell signatures linked to infection and host-dependent differences in parasite transcriptomes, we profiled samples of unexposed, mock- and *P. vivax*-exposed MPCCs across all timepoints. Integrating the data from 31,767 high-quality human transcriptomes with Seurat revealed 8 human clusters, likely representing diverse hepatic cell states (**Figures 4A, S4A)**. Automated annotation of the clusters using SingleR tool identified clusters H0 to H3 as hepatocytes, exhibiting canonical markers of hepato-specific functions (**Figure S4B**). Albeit in small numbers, the remaining clusters H4 to H7 appear to include other cell types besides hepatocytes, such as endothelial and immune cells, likely present in the primary human liver sample used to generate the MPCC. Notably, the cells in the outlying cluster H8 express classical immune markers such as *CD45*, *CD4* and *CD32*.

**Figure 4.**
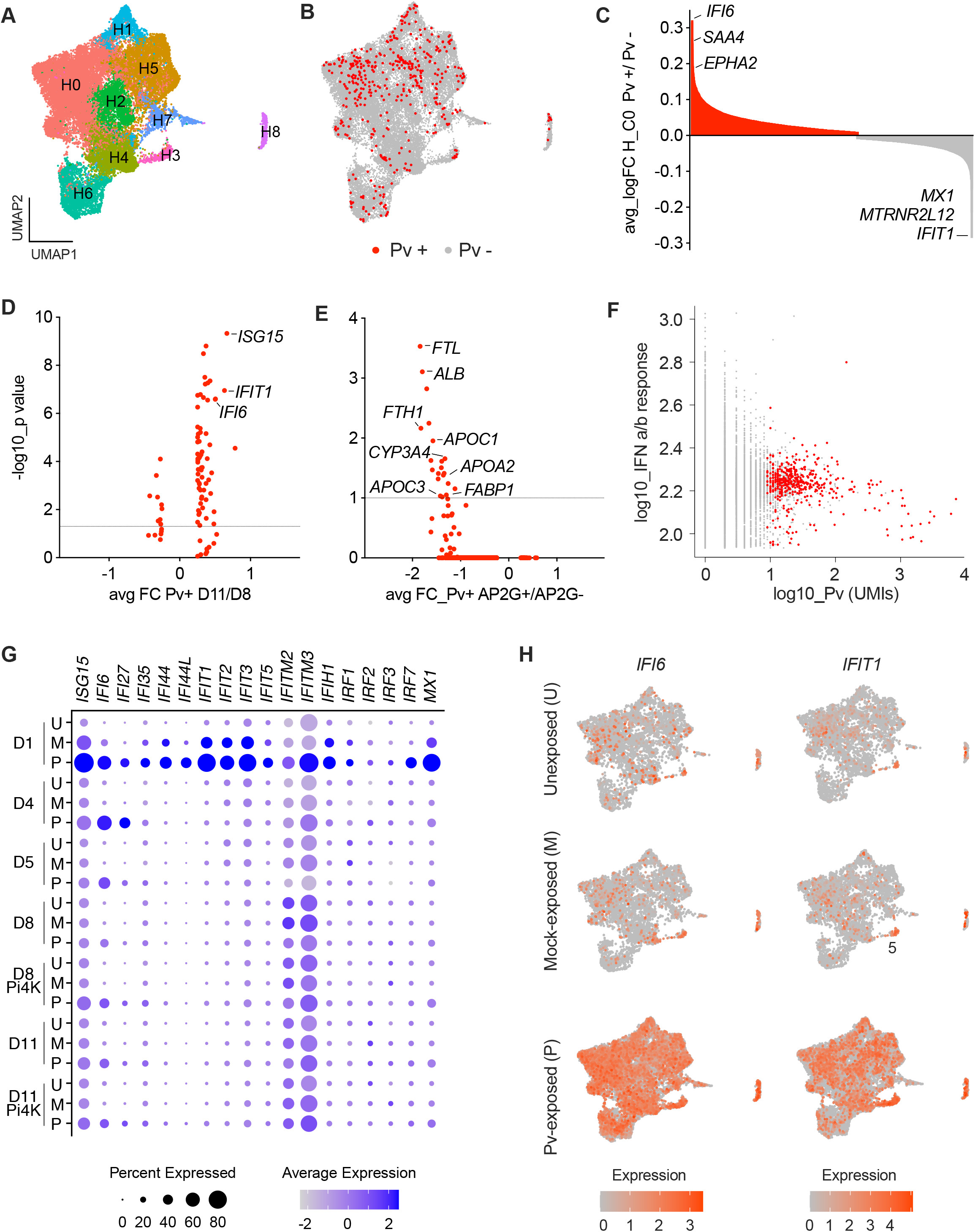
Dual scRNA-seq analysis of *P. vivax* liver-stage infection. **A.** UMAP of 31,767 individual cells colored by cell type cluster using Seurat. Sample identity and cell composition within each human cluster are shown in **Figures S4A-B**. **B.** UMAP as in **A** highlighting the *P. vivax* -positive hepatocytes in red. GSEA analysis in **Figure S4C**. **C.** Waterfall plot comparing *P. vivax* -positive versus -negative in cluster H0 (adjusted *p* < 0.05). **D-E.** Volcano plots comparing differential expression of *P. vivax* -positive cells on day 11 and day 8 (**D**) and AP2-G positive and negative (**E**). **F.** Scatter plot showing higher interferon (IFN) response in *P. vivax* negative cells (grey) acrross all clusters. *P. vivax* -positive hepatocytes are colored in red in **C-F**. **G.** Dot plot showing expression of IFN-related genes throughout infection time course in naïve unexposed (U), mock-exposed (M) and *P. vivax* -exposed (P) samples. **H.** UMAP as in **A** highlighting *IFI6* and *IFIT1* expression levels in U, M and P samples.

*P. vivax-*associated hepatocytes were embedded throughout the cluster landscape, without a clear segregation between infected and non-infected cells (**Figure 4B**). Nevertheless, comparing *P. vivax*-positive and -negative cells in general demonstrated that infected cells are transcriptionally distinct (**Figures 4C, S4C; Table S3**). Gene set enrichment analysis revealed suppression of immune defense and liver-specific functions and increased expression of genes related to membrane organization and modulation by symbiont of host cellular process in *P. vivax*-infected cells (**Figure S4C**). Examples of differentially expressed genes include markers of inflammation and stress response, such as the family of acute-phase serum amyloid A proteins (*SAA4*) and the NFKB inhibitor alpha, caspases and apoptosis regulators (*BIRC3* and *IFI6*), and a glucose transporter (*SLC2A2*) with increased expression in the *P. vivax-* infected population. In contrast, interferon (IFN) regulatory factors (*IRF7*) and IFN induced proteins (*MX1*) are expressed at low levels in *P. vivax*-positive cells. Notably, *EPHA2* was found expressed in higher levels in 50% of the infected cells, consistent with a recent report identifying it as a host factor for human-infecting *P. falciparum* and rodent parasites (Kaushansky et al., 2015). Comparing infected hepatocytes that harbored mid- and late-stage parasites, we found IFN-related genes with significant increased expression in late infection, such as *ISG15*, *IFIT1* and *IFI6,* which might be involved in creating a permissive environment for parasite late-stage development and replication (**Figure 4D; Table S3**).

Given that sexual differentiation and gametocytogenesis in malaria parasites have been linked to host-derived physiological signals (Brancucci et al., 2017), we next compared the transcriptomes of hepatocytes hosting *P. vivax* parasites expressing the transcription factor *AP2-G* versus *AP2-G* negative. Interestingly, we found that hepatocytes bearing *AP2-G* positive parasites exhibit significant lower expression of genes encoding proteins implicated in intracellular iron storage (*FTL* and *FTH1*) and lipid metabolism (*APOA2, APOC1, APOC3, FABP1, CYP3A4*). Genes encoding albumin (*ALB* and *TTR*) and some mitochondrial proteins are also differentially expressed (**Figure 4E; Table S3**). These observations suggest that deficiency in specific intracellular metabolites (possibly iron or lipids) could serve as trigger for *AP2-G* induction and pre-erythrocytic sexual commitment.

### Interferon responses in uninfected bystander hepatocytes

Taking advantage of our single-cell approach, we also investigated the impact of *P. vivax* infection in the neighbor hepatocytes. Comparing the transcriptomes of cells exposed to *P. vivax* to mock-exposed or naïve hepatocytes, we detected a widespread induction of the alpha/beta IFN response, mainly in the *P. vivax-* exposed uninfected cells (**Figures 4F-H**). Day 1 samples stand out given that the induction of multiple IFN-responsive genes and gene families, including *IRF7*, *IFI6*, *IFI27, IFIT1-5, IFI44* and *IFIH1* is observed in more than 40% of the hepatocytes (**Figure 4F**). Transcripts of effector IFN-induced transmembrane proteins *IFITM2* and *IFITM3* are increased throughout the infection time course (**Figure 4F**). The observed temporal order and gene composition of the IFN response to *P. vivax* is remarkably similar to that of hepatitis C virus (HCV) infection (Sheahan et al., 2014), suggesting a common hepatocyte defense mechanism in response to hepatotropic pathogens.

To validate our findings with an independent approach, we performed immunofluorescence analysis of IFITM3 in *P. vivax*-infected MPCCs at a late infection time-point. We found that IFITM3 protein localizes in the bile canaliculi and tight junctions and confirmed the upregulation of IFITM3 in the uninfected bystander cells, with higher expression detected in the vicinity of infected hepatocytes (**Figures 5A-B**). IFITM3 has recently been shown to enhance the trafficking of virus particles to lysosomes (Spence et al., 2019), and thus could be involved in targeting parasites for degradation. Based on its localization, IFITM3 could also be involved in gap junction communication and propagation of host responses from infected cells to uninfected neighbors (Luther et al., 2020; Patel et al., 2009).

**Figure 5.**
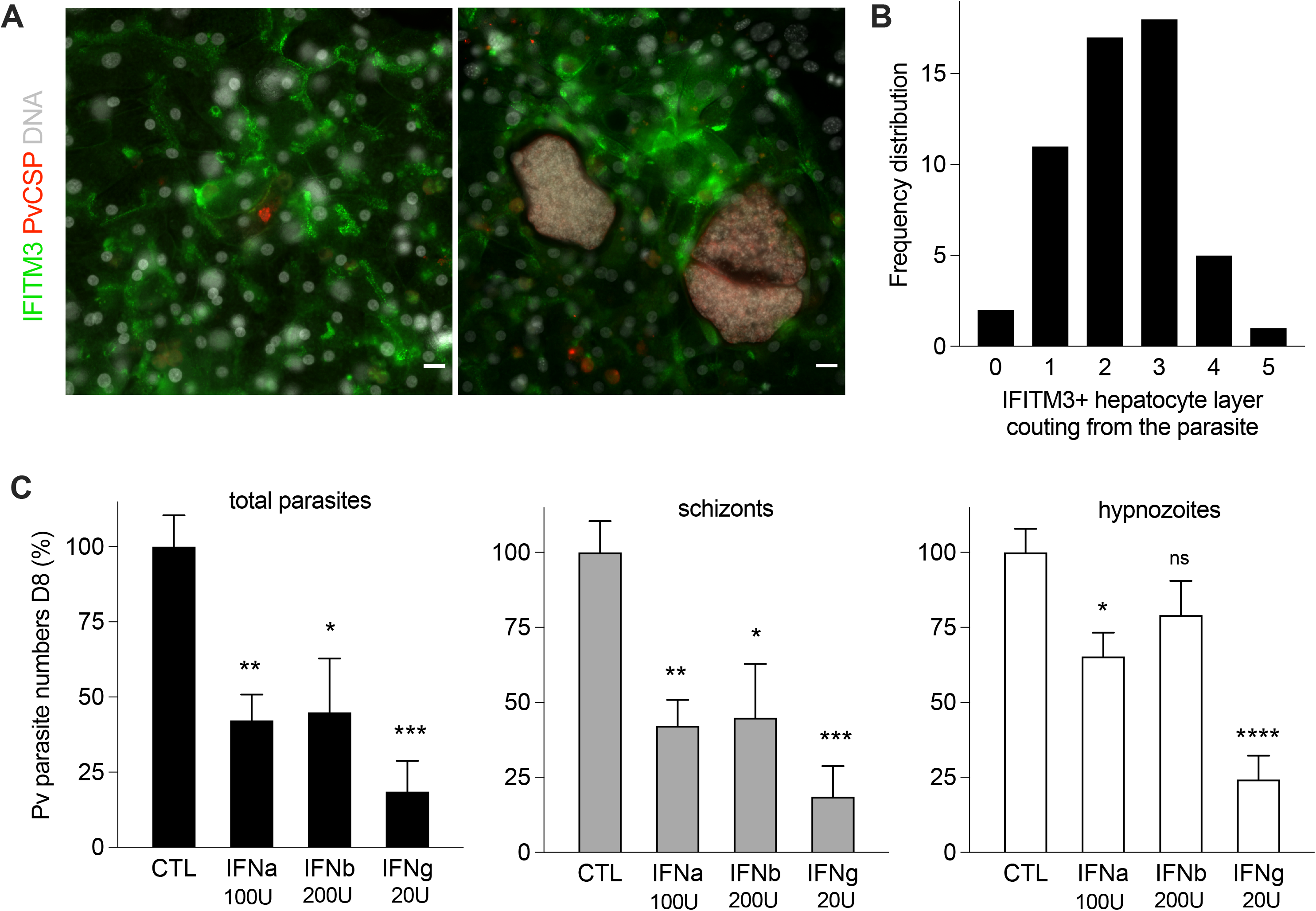
IFN responses in *P. vivax* -infected and bystander hepatocytes. **A.** Expression of IFITM3 protein (green) in *P. vivax-* infected cultures at day 10. Representative images of small (left) and large (right) parasites. Scale bars, 20 μm. **B.** Quantification of IFITM3 positive hepatocytes relative to the infected cell. **C.** IFN treatment (days 5-8) of *P. vivax* infected cultures. Bar plots show quantification of parasite numbers at day 8 (mean ± SEM; *n* = 5-6 wells pooled from 2 independent infections; 1-way ANOVA test: *, *p* < 0.05; **, *p* < 0.01; ***, *p* < 0.001; ****, *p* < 0.0001; *ns*, not significant). Parasite size distribution is given in **Figure S5**.

### Host interferon responses control *P. vivax* infection

Given the induced innate immune responses observed during early *P. vivax* infection, we hypothesized that the activation of this pathway could be associated with the significant drop in parasite numbers observed from day 1 to 11 (**Figure S1A**). To assess the impact of activated IFN signaling on the progression of *P. vivax* infection in MPCCs, we treated infected cultures with IFN alpha (IFNa) and beta (IFNb) cytokines. As control, we used IFN gamma (IFNg), which is known to reduce *P. vivax* infection (Boonhok et al., 2016; Ferreira et al., 1986). While no effect was detected in terms of the kinetics or frequency of parasite development (**Figure S5**), the number of both *P. vivax* schizonts and hypnozoites were significantly reduced upon treatment with IFNa (**Figure 5C**). IFNb appeared to have a stronger effect on schizont stages, by contrast IFNg treatment revealed a potent anti-hypnozoite activity, which is shown here for the first time. Altogether, the results suggest that successful *P. vivax* infection likely involves robust mechanisms for subverting host IFNa/b responses, which might be long-lasting in the hepatocytes that harbor dormant hypnozoites.

## DISCUSSION

Here we comprehensively surveyed the molecular composition of *P. vivax* parasites in distinct liver-stages, as well as their host or neighboring hepatocytes, throughout the course of infection, providing the first single-cell liver atlas of relapsing human malaria. Leveraging the portability of two established platforms (MPCC and Seq-Well) to culture and collect individual human hepatocytes infected in an endemic location, we performed dual single-cell sequencing analysis and further developed a method to selectively enhance capture of parasite transcripts. We demonstrated the robustness and utility of this approach by sequencing 1,991 *P. vivax* parasites and 31,767 hepatocyte transcriptomes and provided validations and mechanistic insights for newly identified gene signatures and parasite-host interactions.

Dual transcriptional profiling of *P. vivax* infection revealed host- and stage-dependent gene expression patterns in both parasites and hepatocytes. Our data is consistent with the following model: upon invasion, a subset of *P. vivax* sporozoites in cells with reduced hepatic metabolism in will activate *AP2-G*. Of those, a fraction will remain dormant, while the remaining portion will activate the schizogony program. Thus, at the point of egress, two populations of hepatic *P. vivax* merozoites will emerge from the liver: the asexual population developing into blood-stages, and the sexually committed population developing into gametocytes for mosquito transmission. This model, while distinct from the existing understanding of the *Plasmodium* life cycle, is in agreement with historical observations of rapid *P. vivax* transmission that occur before the onset of symptoms (Baker, 2010), and is consistent with the high abundance of gametocyte transcripts detected in blood collected from malaria patients (Adapa et al., 2019; Kim et al., 2019). In our model, parasites that do not activate schizogony, the hypnozoites, remain in a dormant state via transcriptional/translational repression mechanisms and rely on proteolytic activity to sustain viability. Bypassing the need for an asexual replication phase might represent an evolutionary imperative to preserve genome integrity, given the prolonged period between infection and transmission. From a clinical perspective, this model would advocate for the development of a novel “wake and kill” clinical approach; namely, to leverage drugs that enhance gametocyte commitment as a way to reduce the hypnozoite reservoir in the liver. Parasite-specific protease inhibitors could also be screened for anti-relapsing activity.

Additionally, our work reveals dysregulation of IFN and inflammatory signaling pathways in *P. vivax*-infected hepatocytes. The reduced expression of IFN-responsive genes in infected cells suggests a protection mechanism for the parasite from host cell-mediated killing. In fact, induction treatments with IFN in our immune-cell free microliver system demonstrated anti-parasite activity against both schizonts and hypnozoites. Downregulation of IFN signaling may also contribute to under-activation of adaptative immune responses by antigen presenting cells, leading to parasite persistence in the organism. However, the mechanisms by which IFN suppression is achieved in infected cells or the impact of host genetics and host-dependent IFN responses, key determinants of clinical outcome in HCV infections (Sheahan et al., 2014), remain unknown. In contrast, we described upregulation of innate immune response pathways in uninfected bystander cells, similar to those observed in viral infections that help control spreading of the virus to neighboring cells (Kotliar et al., 2020; Sheahan et al., 2014). In the context of malaria, this phenomenon might prime or protect cells from a secondary infection, as suggested by the inhibition of malaria re-infection by IFN in a rodent model of *Plasmodium* (Liehl et al., 2013, 2015). Future studies employing spatial single-cell transcriptomics could reveal *P. vivax*-specific gene signatures and spatially heterogeneous responses in bystander cells.

In summary, this study presents the first transcriptional description of individual *P. vivax* liver-stages, their host cells and uninfected bystander cells over the course of infection in human hepatocytes. Leveraging multiple single-cell technologies (sequencing, *in situ* hybridization, and immunofluorescence), we reveal earlier-than expected sexual commitment during *P. vivax* liver-stage development in both replicative and non-replicative parasites, and a dominant innate immune response that exhibits distinct signatures in infected and uninfected bystander hepatocytes. Taken together, these clinically relevant insights provide a framework for characterizing host-parasite interactions in *P. vivax* infections. We expect the methods described here to be applicable for profiling not only other *Plasmodium* parasites at various stages of the life cycle, but also other intracellular pathogens where low abundance of transcripts or host contamination make it difficult to perform single-cell studies.

## ACKNOWLEDGMENTS

We are grateful to Sabrina Hawthorne and Alexei Stortchevoi for help with the targeted Seq-Well; Dan Neafsey, Bronwyn MacInnis, and Sandra Pellegrini for insightful discussions. This work was supported by the Bill & Melinda Gates Foundation (OPP1023607), the National Institute of Health (5U19AI110818-07, 5U24AI118672, P30-CA14051), a Sloan Fellowship in Chemistry (A.K.S.), and a Broad Institute BN10 grant. S.N.B. is a Howard Hughes Medical Institute investigator.

## MATERIALS & METHODS

### Fibroblasts and primary cell cultures

J2-3T3 male murine embryonic fibroblasts (gift of Howard Green, Harvard Medical School) were cultured at < 20 passages in medium comprising of Dulbecco’s Modified Eagle Medium (DMEM, Corning), 10% (v/v) bovine serum (Thermo Fisher Scientific), and 100 mg/mL penicillin-streptomycin (Corning) and were kept at 37°C in a 5% CO2 environment.

Cryopreserved primary human hepatocytes were purchased from BioIVT, a vendor permitted to sell products derived from human organs procured in the United States of America by federally designated Organ Procurement Organizations. Human hepatocytes (male donor, age 57) were maintained in DMEM with 10% fetal bovine serum (FBS, GIBCO), 1% ITS (insulin/transferrin/selenous acid and linoleic acid, BD Biosciences), 7 ng/mL glucagon (Sigma-Aldrich), 40 ng/mL dexamethasone (Sigma-Aldrich), 15 mM HEPES (GIBCO), and 100 mg/mL penicillin-streptomycin (Corning). Hepatocyte cultures were kept at 37°C in a 5% CO2 environment.

### *P. vivax* parasites

*Anopheles dirus* mosquitoes were fed on blood collected from symptomatic patients attending malaria clinics in Tak, Songkla, and Ubon-Ratchathani Provinces in Thailand, confirmed positive for only *P. vivax* via microscopy and RT-PCR. Briefly, *P. vivax* infected blood was drawn into heparinized tubes and kept at 37°C until processing. Infected blood was washed once with RPMI 1640 incomplete medium. Packed infected blood was resuspended in warm non-heat inactivated naive human AB serum for a final hematocrit of 50%. Resuspended blood was fed to laboratory reared female *Anopheles dirus* mosquitoes for 30 minutes via an artificial membrane attached to a water-jacketed glass feeder kept at 37°C. Engorged mosquitoes were kept on 10% sugar at 26°C under 80% humidity at the designated insectary at the Mahidol Vivax Research Unit. Sporozoites were dissected from the salivary glands of infected mosquitoes 14-21 days after blood feeding and pooled in DMEM supplemented with 200 mg/mL penicillin-streptomycin.

### MPCCs and *P. vivax* infection

Primary human hepatocytes were seeded on collagen-micropatterned 96-well plates and surrounded with murine embryonic fibroblasts 3T3-J2s as detailed previously (March et al., 2015). For the scRNA-seq analysis, MPCCs were established using 3T3-J2s expressing an inducible apoptosis switch (Chen et al., 2020). MPCCs were infected with fresh sporozoites obtained through dissection of *P. vivax*-infected mosquitoes. For mock samples, MPCCs were exposed to material from non-infected mosquito salivary glands (matched number of dissected mosquitoes). Unexposed samples received the vehicle. For scRNA-seq, *P. vivax*-infected cultures were collected from duplicate wells, whereas mock and unexposed samples represent single wells for each time-point. For IFN treatments, MPCCs were infected in triplicate wells.

### Drug treatments

To obtain hypnozoite-enriched samples, *P. vivax*-infected MPCCs were dosed with a schizont-specific drug Pi4K inhibitor (MMV390048, 1 μM) for 3 days starting on day 5 after infection. Similar dosing schedule was used for treatments with IFN alpha 2a (11100-1), beta (8499-IF-010) and gamma (285-IF-100) (all purchased from R&D Systems). Cytokine units tested are given in the figures. E64 treatment started at 4h post-infection after media washing the cultures.

### Seq-Well and hybrid capture

For the scRNA-seq analysis, an updated Seq-Well protocol including a second-strand synthesis step was employed (Hughes et al., 2020). Briefly, MPCCs were first treated with AP20187 (0.5 μM) for 30 minutes at 37°C to partially deplete the fibroblast cells via apoptosis. After washing, the cultures were dissociated by Trypsin (0.25%) 5-minute treatment at 37°C. A suspension with 10-15,000 cells was then loaded onto a functionalized-polydimethylsiloxane array preloaded with uniquely barcoded mRNA capture beads. After cells had settled into wells, the array was sealed with a hydroxylated polycarbonate membrane with a pore size of 10 nm, facilitating buffer exchange while permitting cell lysis, mRNA transcript hybridization to beads, and bead removal before proceeding with reverse transcription. The obtained bead-bound cDNA product then underwent Exonuclease I treatment to remove excess primer before proceeding with second-strand synthesis and PCR amplification.

To capture parasite reads from Seq-Well, full length cDNAs were amplified an additional 5 cycles using Kapa HiFi polymerase including a 3-minute extension time to increase concentration for capture. 200-300 ng of cDNA was concentrated using a speedvac, reconstituted in 3.4 μL water and hybridized for 32 hours as previously described (Gural et al., 2018). 10 μL of capture material was amplified for 15 cycles following the standard protocol. Captured cDNA was then prepared into Illumina libraries using NexteraXT (Illumina). Final libraries were quality controlled using Fragment Analyzer (Agilent) and qPCR prior to Illumina sequencing (NextSeq500).

### Quantitative RT-PCR

Total RNA from pooled triplicate wells of *P. vivax*-infected MPCCs was extracted with TRIzol (Thermo Fisher), DNAse treated and purified using the RNeasy MinElute Cleanup Kit (Qiagen). cDNA synthesis was performed using SuperScript II (Thermo Fisher) and RT-PCR was carried out using PowerUp SYBR Green Master Mix (Applied Biosystems) in a Roche Light Cycler 480 Real-Time PCR Detection System according to the manufacturer’s instructions. The primers used are listed in **Table S4**. Relative gene expression was calculated with the delta-delta Ct method, using PVP01_1213400 as housekeeping gene.

### Immunofluorescence analysis

*P. vivax*-infected MPCCs were fixed in ice-cold methanol or 4% paraformaldehyde (PFA), washed in phosphate-buffered saline (PBS) and stored at 4°C. Parasites were detected using *P. vivax*-specific antibodies (PvUIS4, PvBip and PvCSP) on methanol fixed cells as described in (Gural et al., 2018). For IFITM3 staining, PFA-fixed cells were permeabilized with 0.2% TritonX100 for 10 minutes at room temperature, washed in PBS and blocked with 2% bovine serum albumin (BSA) in PBS for 30 minutes at room temperature. IFITM3 rabbit monoclonal antibody (59212, Cell Signaling) was incubated overnight at 4°C (1:100). Alexa-conjugated 488 secondary anti-rabbit antibody (1:1000) was incubated for 1 hour at room temperature, followed by nuclear staining with Hoechst. Images were captured on a Nikon Eclipse Ti or Zen-ApoTome inverted wide-field microscopes using 20x objectives. To quantify parasite size, the area of parasite defined by the PvUIS4 staining was measured using NIS-Elements Microscope Imaging Software and automatically converted to equivalent diameter.

### Fluorescence in situ hybridization

*P. vivax*-infected MPCCs were fixed in 3.7% PFA for 10 minutes at room temperature, washed in PBS, immersed in 70% ethanol and stored at 4°C. Custom labelled probes set specific to *Pv*18S rRNA (FAM dye) and *PvGEXP5* (Quasar 670 dye) purchased from Stellaris were hybridized overnight in the dark at 37°C following the manufacturer’s instructions. After nuclear staining and washing, cells were imaged in a Nikon Eclipse Ti fluorescence microscope as described above.

### Sample Sizes and Statistical Analysis

*n* represents the number wells from each plate as described in the figure legends. Exception for Figures 2C and 2E, where *n* represents 2 independent infections. Methods used for computing statistical significance are indicated in figure legends. Statistical significance was considered for p values below 0.05. Data was analyzed using GraphPad Prism Software.

### Bulk RNA-seq analysis

fastq files were mapped using STAR v. 2.5.3a (Dobin et al., 2013) against PVP01 *P. vivax* v1 genome assembly and annotation, and quantitated by RSEM v. 1.3.0 (Li and Dewey, 2011). Differential expression analysis was performed in DESeq2 on protein-coding genes (Love et al., 2014).

### Tri-genome mapping target generation

Chromosomes and contigs from human (hg19), murine (mm10) [same ENSEMBL releases as in (Macosko et al., 2015)] and *P. vivax* genome (PvP01 v1 release) were renamed with a species-specific prefix and concatenated into a fasta file. Corresponding gtf files were adapted to match prefixed chromosome names, and genes names were adjusted as an ENSEMBLID_SPECIES_GeneSymbol concatenated string.

### scRNA-seq processing

For each sample, fastq files originating from multiple sequencing runs were concatenated and processed using an analytical pipeline derived from the DropSeq pipeline v. 1.12, as described in (Gierahn et al., 2017) (https://github.com/broadinstitute/Drop-seq). Briefly, reads were converted to a bam file using picard v. 2.9.0-1-gf5b9f50-SNAPSHOT, tagged with cell and transcript barcodes, and subsequently sequencing adapters and polyadenosine tracts were trimmed. Upon regenerating fastq files, reads were aligned with STAR v. 2.5.3a (Dobin et al., 2013) against the aforementioned tri-genome reference. Genomic features of the aligned reads were annotated using the combined species-gene symbol nomenclature including gene and exon of origin when relevant. Bead synthesis errors were assessed and when possible, altered unique molecular identifiers (UMIs) were repaired. Cell barcode abundance was tallied, and gene expression was called for the top 10,000 cell barcodes. Count matrices of genes x cells were imported in the R v. 3.6 statistical environment and Seurat v.3 was used as the primary analytical package (Satija et al., 2015). Count matrices were merged into a single Seurat object, which was split by the sample of origin. Human, murine and *P. vivax* genes and transcripts counts were tallied for each cell barcode.

### scRNA-seq analysis of human transcriptomes

Log10 ratios of human and murine-mapped transcripts per cell barcodes were calculated, and only cell barcodes with a ratio greater than 0 were retained. Dataset integration features were selected using Seurat’s SelectIntegrationFeatures function, picking 3,000 genes and subsequently used to prepare the dataset for SCT-based integration PrepSCTIntegration). Integration anchors were identified using the FindIntegrationAnchor function, with parameters “SCT” as a normalization method, the aforementioned list of 3,000 genes as anchor features, 30 PCA dimensions, k.anchor=10 and using the first sample as reference (Butler et al., 2018). This integrated Seurat object was further filtered and cell barcodes displaying a human/murine coverage greater than 10-fold were retained for downstream analyses, resulting in 31,767 cells remaining. The integrated object was scaled and centered using a linear model, with scale.max=10, block.size =1,000 and min.cells.to.block=3,000. Principal component analysis was performed using RunPCA, retaining the top 30 components. The number of principal components retained was picked based on the inspection of the “elbow plot” and JackStraw procedure (n=100 replicates, 50 PCs) in Seurat. A UMAP embedding was calculated on the top 30 principal components using the RunUMAP function with flags n.neighbors=30, metric=cosine, learning.rate=1, min.dist=0.3, spread=1, set.op.mix.ratio=1, local.connectivity =1, repulsion.strength =1, negative.sample.rate=5, uwot.sgd= FALSE, seed.use: 42, angular.rp.forest: FALSE on the integrated object. Cluster identification was based on the Louvain algorithm for nearest-neighbor identification: the top 30 principal components were used to build the kNN graph, considering 20 nearest neighbors, with prune.snn=0.067, nn.method was an exact RANN (nn.eps=0) and Euclidean distance as the annoy. The resulting graph was partitioned using a shared nearest neighbor (SNN) modularity optimization-based clustering algorithm at resolution 0.8 to identify clusters (with 10 starts and 10 iterations, and standard modularity function). Gene and transcript coverage plots per cell barcodes were generated on the integrated dataset prior to the final filtering step (for barcodes with a 10-fold enrichment for human transcript). Cluster markers were identified using the FindAllMarkers function with logfc.threshold set to −100 and at least 10% of cells expressing the marker. Differential gene expression signature was queried between *P. vivax*-positive and -negative cells using a Wilcoxon test (implemented in the FindMarker function). Cell barcodes positive for eight or more parasite genes were deemed to be *P. vivax*-positive.

### scRNA-seq analysis of *P. vivax* transcriptomes

The Seq-Well analytical pipeline was run on the samples as described above. For samples for which pre- and post-capture data were available, matching cell barcodes were identified in the corresponding genes x cell matrices and UMI counts were summed over pre- and post-capture libraries. The resulting matrices were merged into a Seurat object (2,171 cells, 85,276 genes). Only *P. vivax* genes were retained for subsequent analysis (n=6,478). Samples which had less than 20 cells were excluded from the analysis. The resulting object (n=1,991 cells) was split by dataset. Raw counts for each sample were exported into individual text files and processed with Scanorama (v. 1.5, under Python 3.6 environment) for normalization, batch correction and integration (Hie et al., 2019). The top 19 integration dimensions were imported back into Seurat and used for data embedding using UMAP, as well as cluster elicitation via the Louvain algorithm (similar to the procedure ran for the human compartment described above). Cluster-specific gene markers were identified using a Wilcoxon test. Subclustering was performed by extracting Scanorama dimensions for the cells of interest, followed by UMAP and Louvain-based cluster identification. Heatmaps were generated in Seurat using hclust-based hierarchical clustering of the scaled data with a centroid distance metric (UPMGC-equivalent), followed by manual editing.

## Notes

### Competing Interest Statement

A.K.S. reports compensation for consulting and/or SAB membership from Merck, Honeycomb Biotechnologies, Cellarity, Repertoire Immune Medicines, Hovione, Ochre Bio, Third Rock Ventures, Relation Therapeutics, and Dahlia Biosciences.

