## Supplemental Figures for "Gene signatures and host-parasite interactions revealed by dual single-cell profiling of *Plasmodium vivax* liver infection"

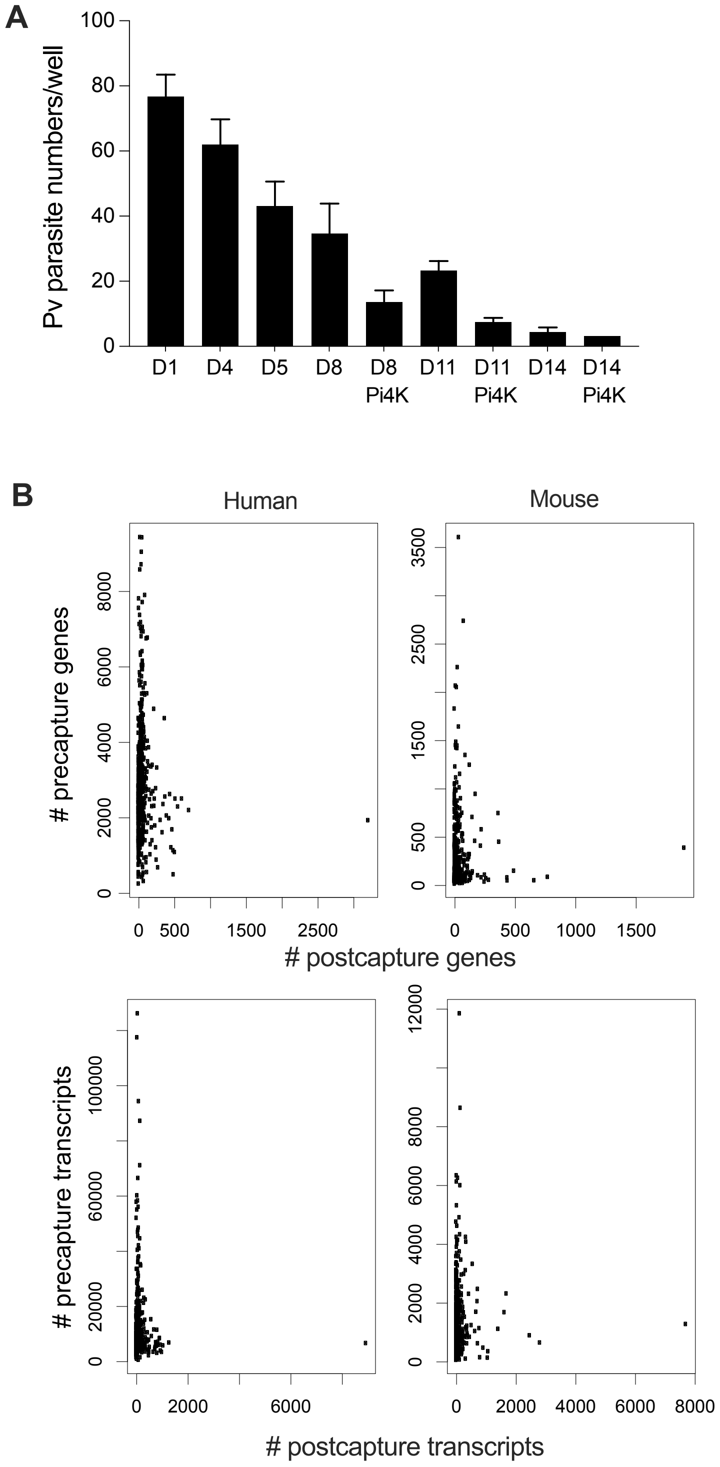


**Figure S1. A.** Quantification of parasite numbers (mean ± SEM; $n$ = 3-8 wells pooled from 2 independent infections used for scRNA-seq). **B.** Pairwise comparisions of the gene and trancript numbers in pre- and post-capture conditions. Human (left) and mouse (right) species shown separately.

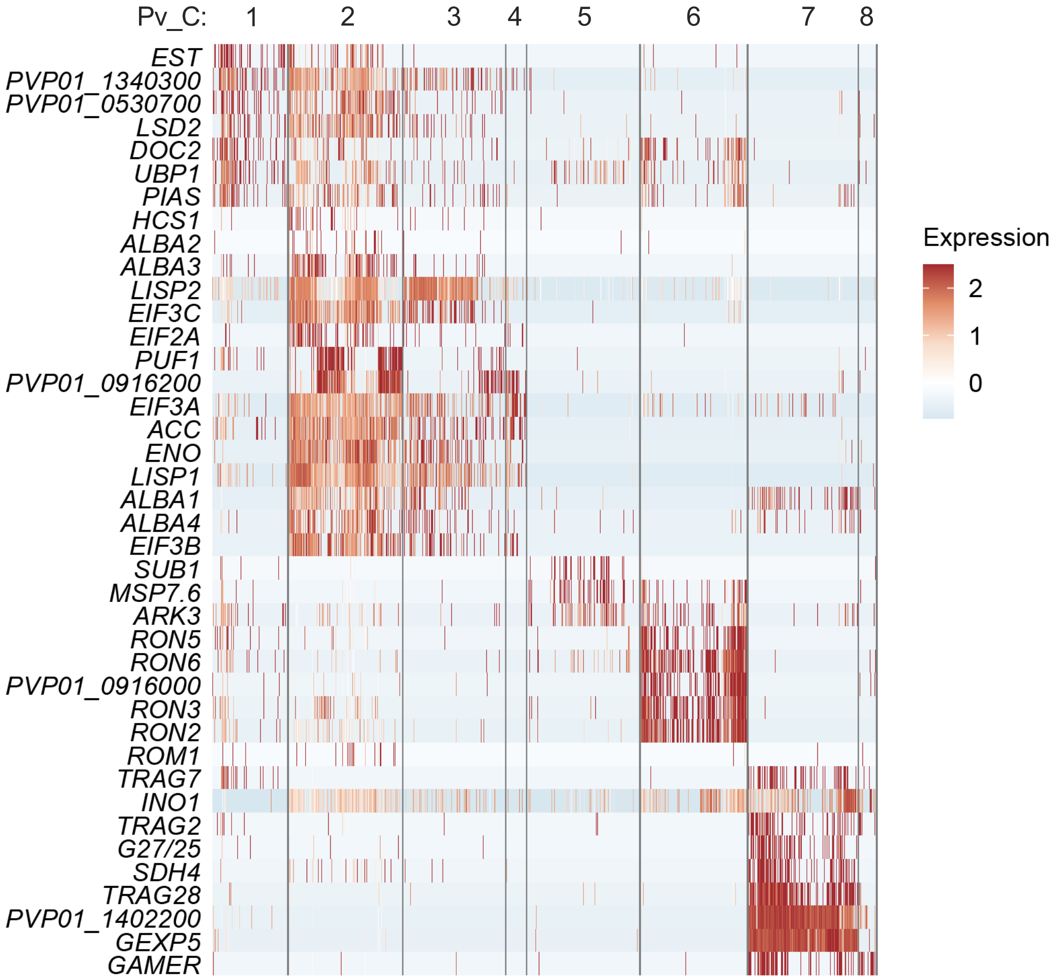


**Figure S2.** Heatmap showing *P. vivax* cluster-specific marker genes. 100 parasites are shown, except clusters Pv_C4 and Pv_C8 that contain fewer cells. Each bar represents a single parasite.

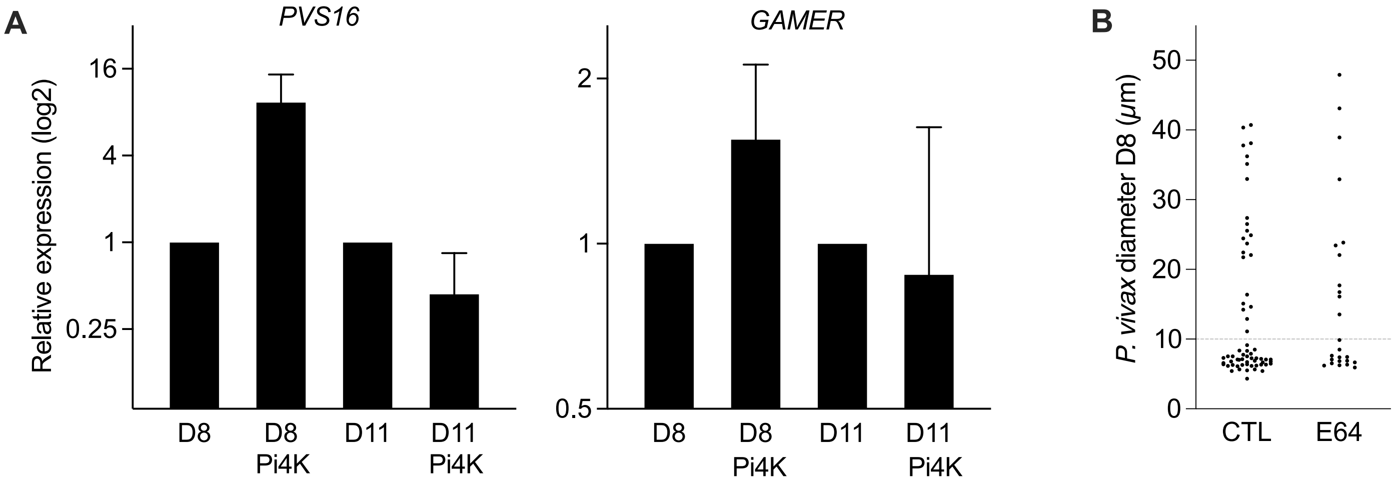


**Figure S3. A.** Quantitative RT-PCR analysis of gametocyte markers (*PVS16* and *GAMER*) in Pi4K treated bulk samples on days 8 and 11. Non-treated samples were set to 1 (mean ± SEM; *n* = 2 independent infections). **B.** Scatter plot showing the parasite size distribution in MPCC cultures treated with protease inhibitor E64 (1 µM) for 8 days. Each dot represents an individual parasite (*n* = 3-5 wells pooled). The traced line indicates the 10 µm cutoff used for hypnozoite staging.

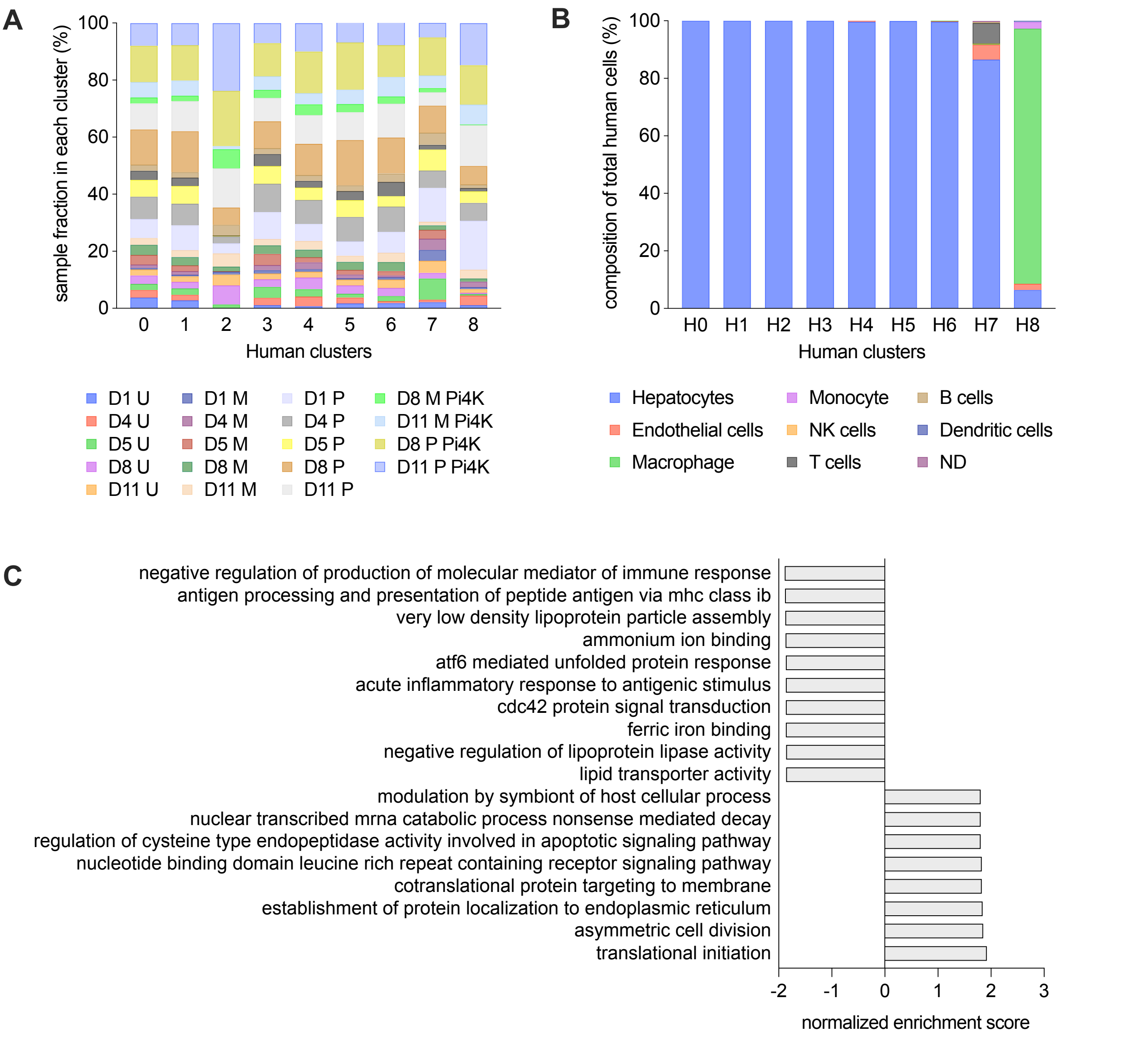


**Figure S4. A-B.** Stacked bar plots showing the sample fraction (**A**) and cell type composition (**B**) whithin each human cluster. U, unexposed; M, mock-exposed; P, *P. vivax*-exposed; ND, not determined. **C.** Gene set enrichment analysis of differentially expressed genes in *P. vivax*-positive versus -negative hepatocytes (H0 cluster). Bars show the top 10 terms for the negative score (full list in **Table S3**). False discovery rate was set at 0.05.

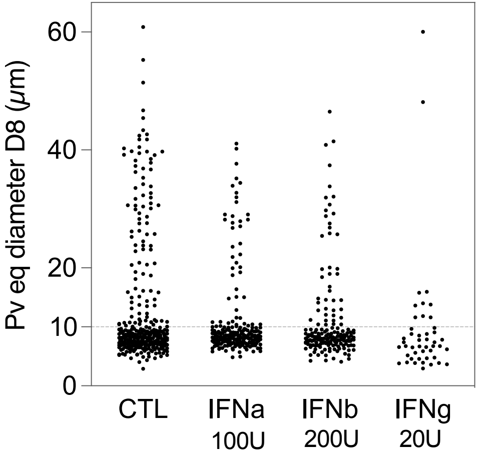


**Figure S5.** Scatter plot showing the parasite size distribution in MPCC cultures treated with interferon alpha (IFNa), beta (IFNb) and gamma (IFNg) for 3 days starting on day 5 after infection. Cultures were fixed and analyzed on day 8. Each dot represents an individual parasite (*n* = 5-6 wells from 2 independent experiments pooled). The traced line indicates the 10 μm cutoff used for hypnozoite staging.
